# A hybrid machine learning and regression method for cell type deconvolution of spatial barcoding-based transcriptomic data

**DOI:** 10.1101/2023.08.24.554722

**Authors:** Yunqing Liu, Ningshan Li, Ji Qi, Gang Xu, Jiayi Zhao, Nating Wang, Xiayuan Huang, Wenhao Jiang, Aurélien Justet, Taylor S. Adams, Robert Homer, Amei Amei, Ivan O. Rosas, Naftali Kaminski, Zuoheng Wang, Xiting Yan

## Abstract

Spatial barcoding-based transcriptomic (ST) data require cell type deconvolution for cellular-level downstream analysis. Here we present SDePER, a hybrid machine learning and regression method, to deconvolve ST data using reference single-cell RNA sequencing (scRNA-seq) data. SDePER uses a machine learning approach to remove the systematic difference between ST and scRNA-seq data (platform effects) explicitly and efficiently to ensure the linear relationship between ST data and cell type-specific expression profile. It also considers sparsity of cell types per capture spot and across-spots spatial correlation in cell type compositions. Based on the estimated cell type proportions, SDePER imputes cell type compositions and gene expression at unmeasured locations in a tissue map with enhanced resolution. Applications to coarse-grained simulated data and four real datasets showed that SDePER achieved more accurate and robust results than existing methods, suggesting the importance of considering platform effects, sparsity and spatial correlation in cell type deconvolution.

## Introduction

Spatial transcriptomic technologies enabled measuring gene expression and physical locations of spots and/or cells simultaneously in intact tissues of various types in an unbiased and high-throughput way [1-4], providing unprecedented information to understand disease associated changes. Specifically, the spatial barcoding-based (ST) technologies, such as Slide-seq[5], HDST[6], ST[4], and 10x Genomics Visium, divide tissue into small capture spots and measure high-throughput gene expression levels unbiasedly for each spot with known physical location [4-10]. Depending on the size of capture spots, the measured expression profile is an average expression profile of cells of unknown types. Therefore, the corresponding data lacks single-cell resolution[11] and requires cell type deconvolution to understand the cell type composition and cell type-specific gene expression in each spot.

One common way to deconvolve ST data is to use cell type-specific expression profile from existing single-cell RNA sequencing (scRNA-seq) data of the same tissue type. Many methods have been developed [12-27], which can be divided into four categories: machine learning-based [20-22], regression-based [23-27], statistical modeling-based [23-27] and data mapping-based methods. Benchmarking studies have been conducted to compare the performance of these methods [28-30].

Despite of the success of current methods, the following three challenges have not been well addressed and more importantly, no method address them simultaneously. First, systematic difference exists between scRNA-seq and ST data[12-15, 22, 24, 25] due to various technical factors, such as differences in protocols, reagents, platforms or simply sequencing depths. This systematic difference, termed as platform effects [12], makes the relationship between ST data and cell type-specific expression profiles from the reference scRNA-seq data non-linear and vary across different technologies. A few statistical model-based methods [12-15] consider the platform effects as multiplicative random or fixed effect. However, these methods were shown in previous benchmarking study [28] to have comparable performance to methods that do not address platform effects, leaving it unclear whether platform effects were adequately addressed. DSTG and some of the data mapping-based methods implicitly addressed platform effects by embedding scRNA-seq or scRNA-seq derived pseudo-spot data and real ST data into a common latent space. Second, among all cell types existed in the tissue, only a few cell types are present in each spot. For example, 38 different cell types were found in the scRNA-seq data of whole lung tissues (the IPF dataset in real data analyses). However, capture spots of the 10x Genomics Visium platform with a size of ∼55μm contained only 2-10 cells per spot, demonstrating a sparse presentation of all cell types existed in the tissue. This sparsity was considered by RCTD, SPOTlight, DestVI and SpatialDWLS but using subjective hard thresholding. Lastly, previous studies [24, 31, 32] have shown that cell type composition of spots that are physically close in the tissue tend to be similar or correlated. Only CARD explicitly considered the across-spot spatial correlation of cell type compositions.

To address all the aforementioned challenges, we propose a two-step hybrid machine learning and regression method, SDePER, that considers platform effects removal, spatial correlation and sparsity (Fig. 1). In the first step, a conditional variational autoencoder (CVAE)[33] is used to adjust the ST and reference scRNA-seq data for platform effects removal. In the second step, a graph Laplacian regularized model (GLRM) is fitted to the adjusted ST data with consideration of the spatial correlation of cell type compositions between neighboring spots and sparsity of present cell types per spot. Based on the estimated cell type compositions, a random walk is performed to impute cell type compositions and gene expression at unmeasured locations in a tissue map with enhanced resolution. We demonstrate the advantage of SDePER through extensive simulations and applications to four real datasets from various tissues, species, and technologies.

**Figure 1.**
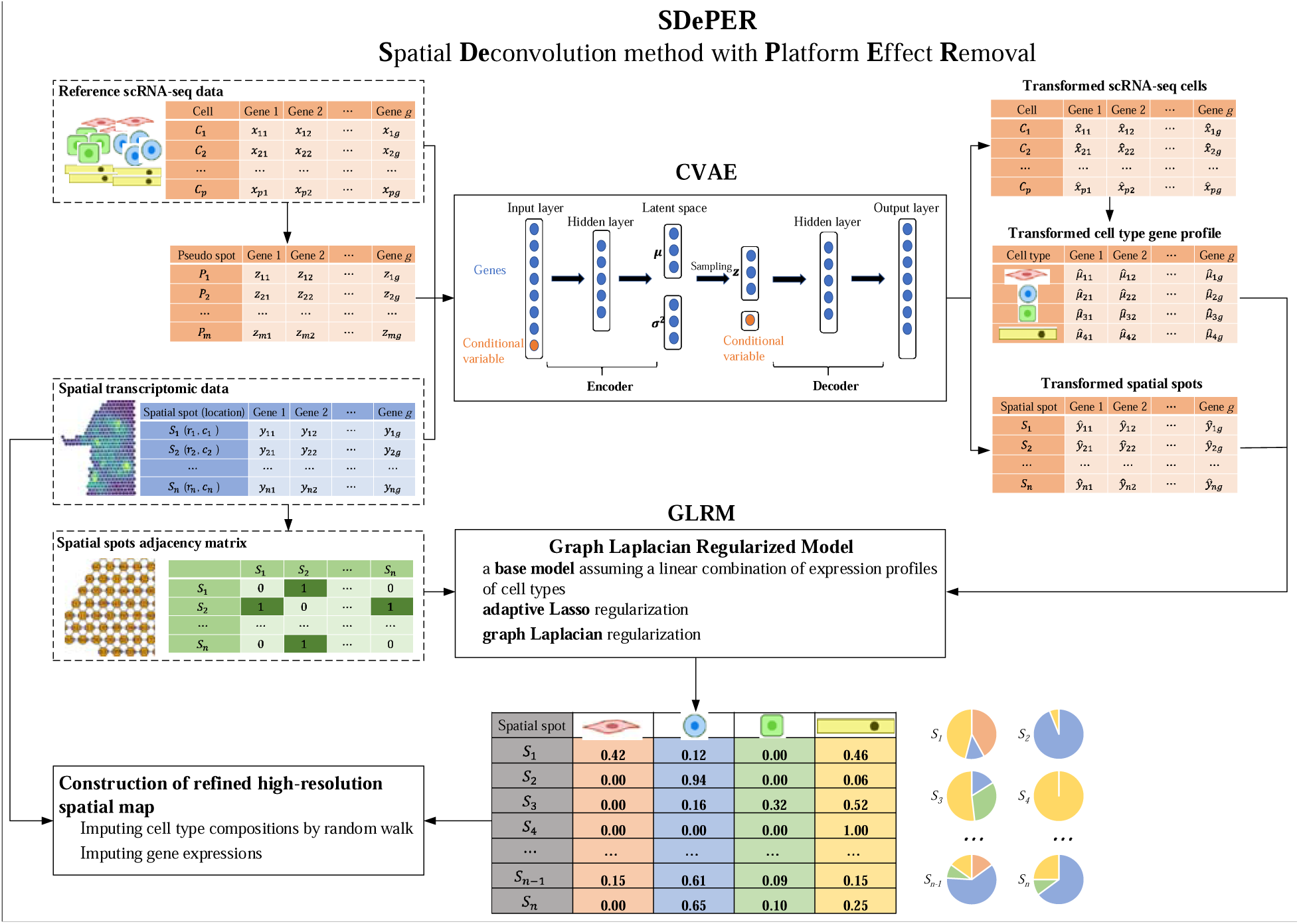
Schematic overview of SDePER. SDePER performs cell type deconvolution of ST data in a two-steps fashion. In the first step, conditional variational autoencoder (CVAE) takes three datasets as input: real ST data, reference scRNA-seq data and pseudo-spot data generated using the reference scRNA-seq data. Using the trained encoder and decoder under the two conditions (ST and scRNA-seq), real ST data is transformed to the same space as scRNA-seq data and pseudo-spot data. The transformed real ST data and cell type-specific expression profiles are then used to fit the graph Laplacian regularized model (GLRM) with penalties for sparsity and across-spot spatial correction in cell type compositions. The estimated cell type compositions from GLRM can be further used to impute for cell type compositions and gene expression at unmeasured locations in the original spatial map to construct new spatial map at arbitrarily higher resolution.

## Results

### SDePER efficiently removes platform effects

We conducted simulations to evaluate the performance of SDePER and compared it with six other deconvolution methods with the best performance based on previous benchmarking studies[11, 28-30]: SpatialDWLS[26], cell2location[15], SPOTlight[25], CARD[24], DestVI[14], and RCTD[12]. ST data was simulated by coarse-graining a real spatial transcriptomic data with single-cell resolution (Fig. 2A) generated using the STARmap technology[34]. The true cell type composition at each simulated spot is calculated and serves as the ground-truth. To demonstrate the impact of platform effects [12] on the method performance, each method was applied using both external and internal reference data, representing situations with and without platform effects. Moreover, to demonstrate the effectiveness of CVAE on removing platform effects, we ran SDePER with the CVAE component deactivated, which was named GLRM.

**Figure 2.**
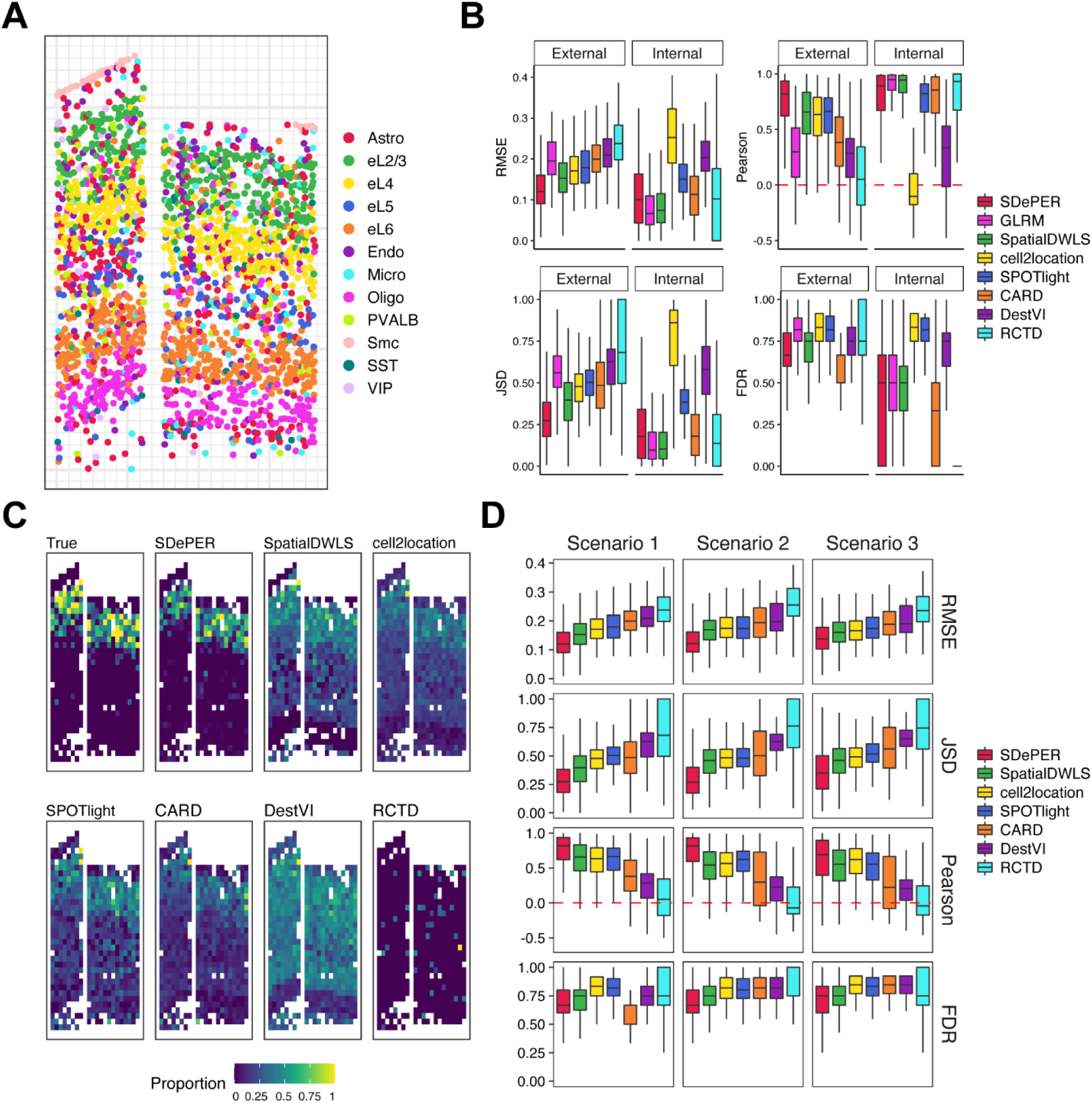
Performance evaluation and comparison using simulation studies. (A) Coarse-graining procedure to simulate ST data with ground-truth. (B) Demonstration of the impact of platform effects on method performance: Boxplots show the median (center line), interquartile range (hinges), and 1.5 times the interquartile (whiskers) of RMSE, JSD, Pearson’s correlation and FDR using external scRNA-seq reference and internal single-cell level spatial reference. (C) The proportion of L2/3 excitatory neurons in the simulated spots. (D) Boxplots show the median (center line), interquartile range (hinges), and 1.5 times the interquartile (whiskers) of RMSE, JSD, Pearson’s correlation and FDR using different scRNA-seq reference: scenario 1: scRNA-seq reference with matched cell type; scenario 2: one missing cell type in scRNA-seq reference; scenario 3: one added irrelevant cell type in scRNA-seq reference.

Performance comparison based on the median RMSE, Pearson correlation and JSD showed that SDePER achieved the highest estimation accuracy regardless of the existence of platform effects (Fig. 2B, S2). Visualization of the ground-truth and estimated proportion of L2/L3 excitatory neurons (Fig. 2C) and other cell types (Fig. S1) using external reference further confirmed the highest accuracy of SDePER results (Pearson correlation=0.839). Furthermore, the accuracy of all methods except cell2location was lower for external reference compared to internal reference, indicating that platform effects have a complicated form that cannot be efficiently addressed using a random effect. SDePER and DestVI had the smallest accuracy difference between internal and external reference, suggesting their best robustness to platform effects (Fig. 2B, S2). Lastly, when using internal reference without platform effects, SDePER had slightly worse performance than GLRM with an increase of 0.021 and 0.063 in the median RMSE and JSD, respectively, and a decrease of 0.040 in the median correlation, indicating the potential noise introduced by the CVAE component. But when platform effects were present (external reference), SDePER had a much better performance than GLRM (29%, 41%, 47%, 6% improvement in RMSE, JSD, Pearson’s correlation, and FDR, respectively) and this increase was much larger than the decrease in performance when using internal reference (Supplementary Table 1). All these demonstrated that SDePER achieved the best performance in both estimating cell type compositions and removing platform effects.

### Robustness of methods to mismatching cell types

To demonstrate the robustness of methods to mismatching cell types between the reference and ST data, we conducted deconvolution under three scenarios representing perfect matching, on missing cell type and one extra cell type in the external reference data compared to the ST data. The performance rankings of all methods were consistent across these three scenarios with SDePER consistently achieving the best accuracy (Fig. 2D). In scenario 1, the improvements in RMSE of SDePER compared to SpatialDWLS, cell2location, SPOTlight, CARD, DestVI, and RCTD were 21%, 29%, 32%, 39%, 42% and 49%, respectively. In scenario 2 and 3, SDePER achieved 24–50% and 26-50% improvement in RMSE, respectively. Compared to scenario 1, SDePER also had an increase of 0.004 and 0.016 in the median RMSE in scenario 2 and 3, respectively. These results showed that SDePER had the best robustness to the mismatching cell types between the ST data and reference scRNA-seq data.

### Mouse olfactory bulb data

To demonstrate the efficacy of SDePER in real data, we first applied SDePER and the other six methods to a ST data of mouse olfactory bulb (MOB) [4] with well-defined anatomic layers organized in a well-characterized spatial architecture. We took an independent scRNA-seq data of the same tissue type profiled using the 10x Genomics Chromium platform as reference data for the deconvolution[35]. Based on the H&E staining, four major tissue layers were identified from inside to outside with each dominantly composed by one cell type: the granule cell layer (GCL), mitral cell layer (MCL), glomerular layer (GL), and olfactory nerve layer (ONL) dominated by GC, M/TC, PGC, and OSNs, respectively (Fig. 3A) [4, 24]. Expression maps of marker genes for these 4 dominant cell types were consistent with the four annotated layers (Fig. 3A, S3).

**Figure 3.**
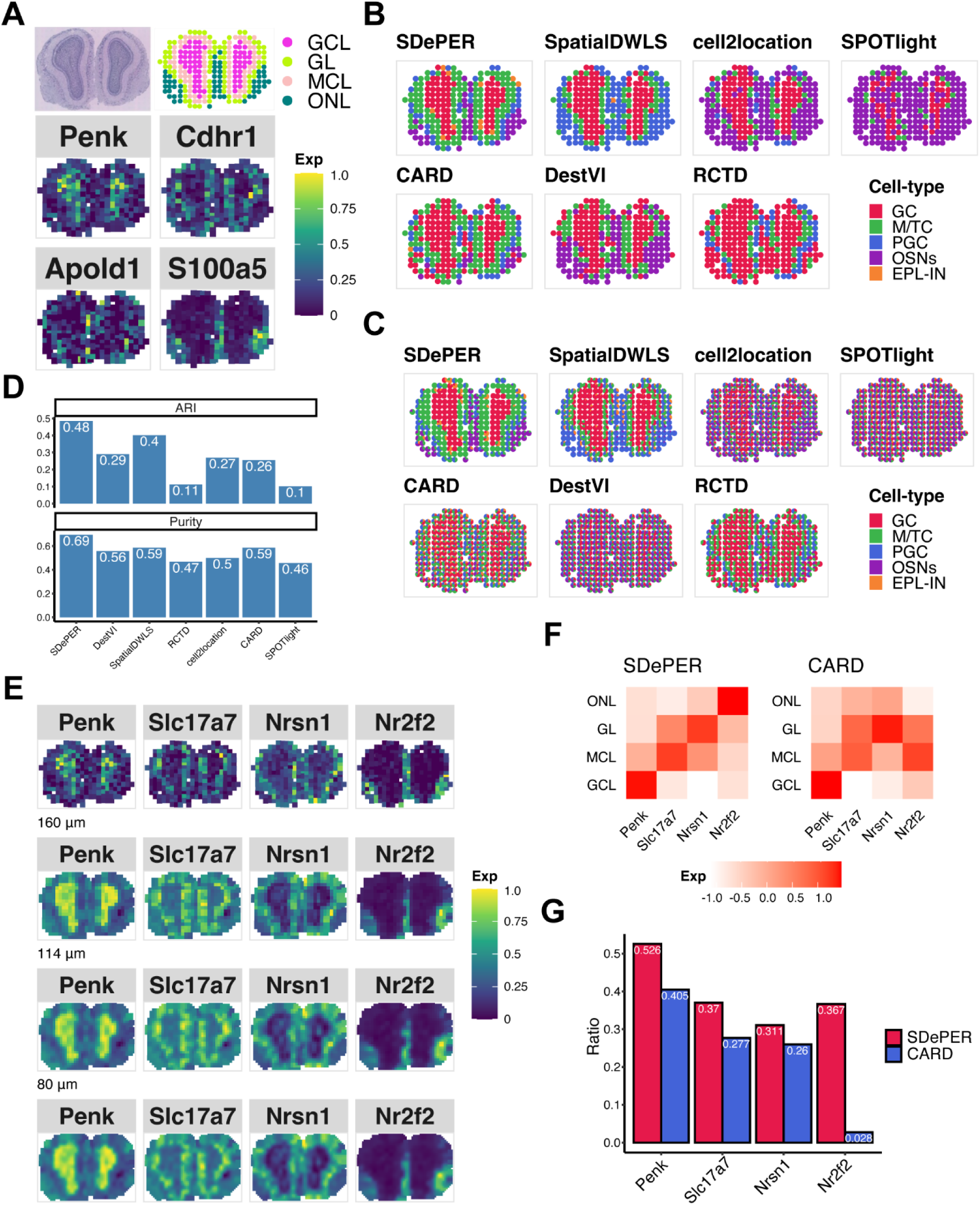
Performance evaluation and comparison using MOB dataset. (A) H&E staining of MOB (top-left), annotated regions (top-right GCL: granule cell layer; MCL: mitral cell layer; GL: glomerular layer; ONL: olfactory nerve layer) and expression pattern of cell-type-specific marker genes (bottom, *Penk* for GC, *Cdhr1* for mitral and tufted cell (M/TC), *Apold1* for periglomerular cell (PGC), and *S100a5* for olfactory sensory neurons (OSNs)). (B) Visualization of inferred dominant cell type in each spot (EPL-IN: external plexiform layer interneuron). (C) Spatial scatter pie chart of estimated cell type composition within each spot. (D) Comparing deconvolution methods using ARI (left) and purity (right). (E) Expression patterns of the corresponding layer-specific marker genes and imputed expression at three different resolution levels: 160μm (about 64% of original size), 114μm (about 32% of original size), 80μm (about 16% of original size). (F) Heatmap showing average imputed expression of region-specific marker genes at 80μm level within each annotated region for SDePER and CARD. (G) Bar plot showing the ratio of average layer-specific marker gene expression in the corresponding layer among all layers.

The H&E staining image-based annotation and expression maps of the four dominant cell type marker genes were considered as ground-truth. The predicted dominant cell type by SDePER showed remarkable similarity with the ground-truth (Fig. 3A, 3B). SpatialDWLS, DestVI and cell2location did not separate ONL and GL. RCTD mislabeled ONL as GCL. CARD showed blurry layer boundaries and did not find ONL accurately. Quantitative assessment of the similarity between the predicted dominant cell type and H&E staining image-based annotated layers using ARI and purity (Fig. 3D) confirmed the best performance of SDePER. In addition, when comparing the predicted dominant cell type (Fig. 3B) to the predicted cell compositions in pie chart (Fig. 3C) for each method, SDePER showed the highest similarity between the two plots indicating less non-specific cell type detection potentially due to its sparsity regularization.

To demonstrate the imputation results, we selected four layer-specific marker genes, one gene for each layer, from the ST data (Fig. 3A). Visualization of their original and imputed expression on original spatial map and three spatial maps with higher resolution (Fig. 3E, S4-5) showed an expression enrichment of each layer marker gene in its corresponding layer. To levels of each layer marker gene in its corresponding layer at 80 *μm* resolution. The average quantitatively assess the expression enrichment, we calculated the average imputed expression imputed expression by SDePER (Fig. 3F) displayed higher diagonal values and lower off-diagonal values, indicating a better separation between different layers based on the imputed expression than CARD. We further calculated the ratio of average expression of each layer-marker gene in its corresponding layer to that across all layers for a quantitative assessment of the imputed expression-based layer separation (Fig. 3G). SDePER achieved a higher ratio for all four layer-marker genes than CARD, demonstrating a higher accuracy in imputed expression.

### Stage III cutaneous malignant melanoma data

The second real data we analyzed investigated the cutaneous malignant melanoma sample from lymph nodes[7]. Manual annotation of the tissue slide using H&E staining and clustering analysis of the ST data (Fig. 4A) identified regions of melanoma, stroma, and lymphoid tissue with expected cell types [7, 32]. For each expected cell type in each region, we selected its marker genes from existing literatures[36], which included *PMEL* for malignant cells in melanoma regions, *COL1A1* for fibroblast in stroma regions, *MS4A1* for B cells and *CD14* for macrophage in lymphoid tissues [36]. The expression map of these marker genes in ST data (Fig. 4A) confirmed the prevalence of fibroblasts in stroma regions, B cells in the right-top lymphoid tissue 1, and macrophages in the lymphoid tissue 2 surrounding the melanoma region. Expression maps of marker genes for other cell types can be found in Fig. S6. We used an independent scRNA-seq data of untreated metastatic melanoma samples from human lymph nodes [37] profiled using the inDrop technology as the reference data for deconvolution.

**Figure 4.**
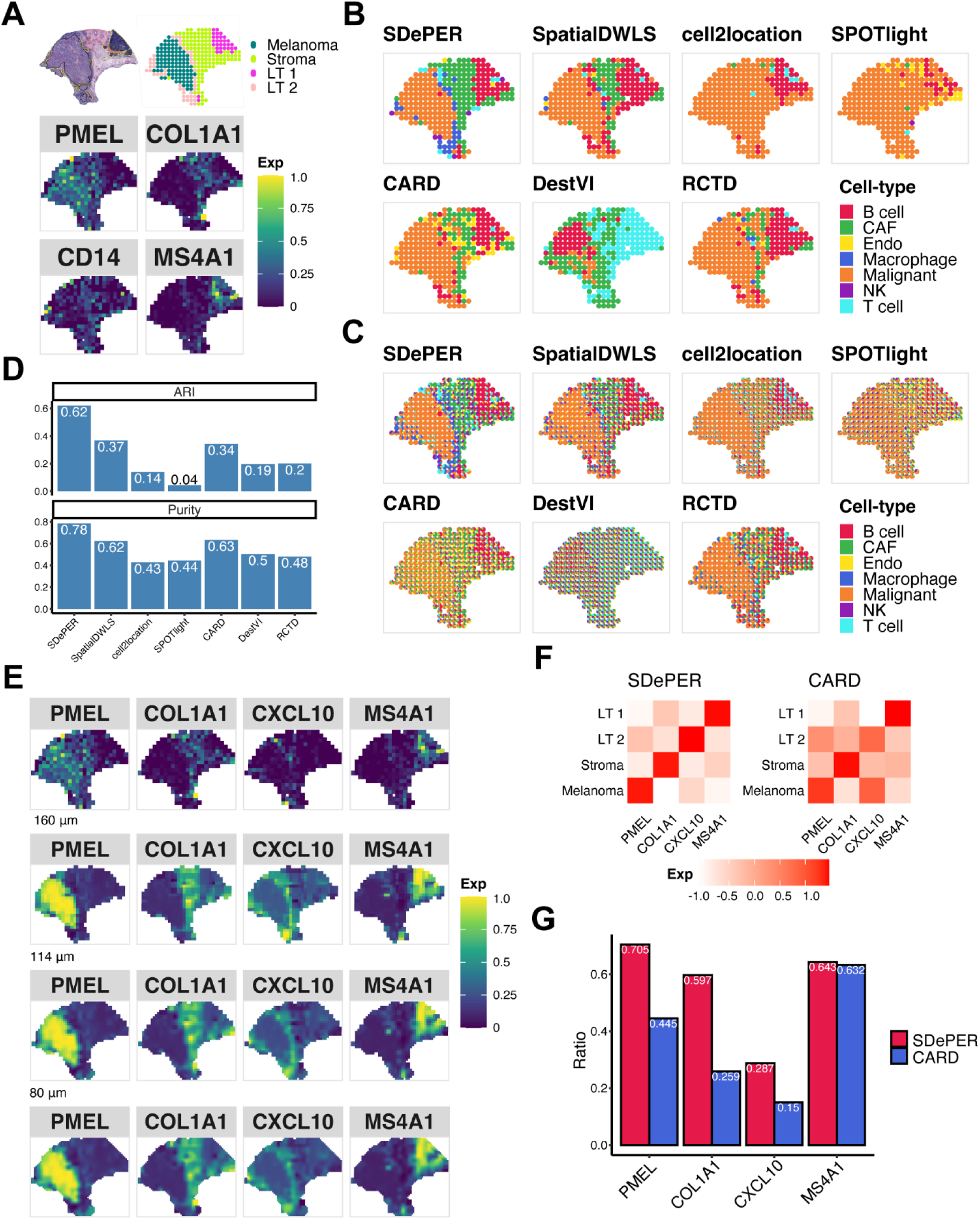
Performance evaluation and comparison using melanoma dataset. (A) H&E staining of melanoma (top-left, melanoma (black), stroma (red), lymphoid tissue (yellow)), annotated regions (top-right, LT: lymphoid tissue) based on BayesSpace and expression pattern of cell-type-specific marker genes (bottom, *PMEL* for malignant melanoma regions, *COL1A1* for fibroblast in stroma regions, *CD14* for macrophage, and *MS4A1* for B cells). (B) Visualization of inferred dominant cell type in each spot (CAF: cancer-associated fibroblasts; Endo: endothelial; NK: natural killer). (C) Spatial scatter pie chart of estimated cell type composition within each spot. (D) Comparing deconvolution methods using ARI and purity. (E) Expression patterns of the corresponding region-specific marker genes and its imputed expression at three different resolution levels: 160μm (about 64% of original size), 114μm (about 32% of original size), 80μm (about 16% of original size). (F) Heatmap showing average imputed expression of region-specific marker genes at 80μm level within each annotated region for SDePER and CARD. (G) Bar plot showing the ratio of average layer-specific marker gene expression in the corresponding layer among all layers.

Like the results of MOB data, the dominant cell type predicted by SDePER highly matched the H&E staining image-based annotation (Fig. 4B). In contrast, other methods failed to identify a clear boundary between regions (Fig. 4B). SDePER also achieved the highest ARI and purity that are 1.68 and 1.24 times higher, respectively, than the second-best method (Fig. 4D). The least non-specific cell type detection by SDePER was again observed (Fig. 4B-C).

Four region-specific marker genes (*PMEL* for melanoma region, *COL1A1* for stroma region, *CXCL10* for lymphoid tissue 2 and *MS4A1* for lymphoid tissue 1) were identified from the ST data (Fig. 4E, Fig. S7-8) to demonstrate the accuracy of imputed expression. As expected, the imputed expression map of each region marker gene by SDePER showed increased expression in the correct region (Fig. 4E). Compared to CARD, the SDePER imputed *CXCL10* expression remarkedly better resembled the lymphoid tissue 2 on the periphery of tumor (Fig. S8). The average imputed expression of each region marker gene by SDePER had a better enrichment for the correct region (Fig. 4F), confirmed by the higher expression ratio of SDePER for each region marker gene (Fig. 4G). All these results suggested that the SDePER imputed gene expression was more accurate.

### HER2-positive breast tumor data

Next, we analyzed the ST data from patients with HER2-positive breast tumor, which consists of various cell types arranged in spatial domains annotated by pathologists [8]. The annotated regions included two cancer regions (cancer in situ and invasive cancer), four named regions (adipose tissue, breast glands, connective tissue, and immune infiltrate), and undetermined regions (Fig. 5A). Although no expected cell type was provided in each region, we do expect the cancer regions to enrich for cancer epithelial cells. An external scRNA-seq dataset from 5 HER2-positive patients was used as the reference data [38].

**Figure 5.**
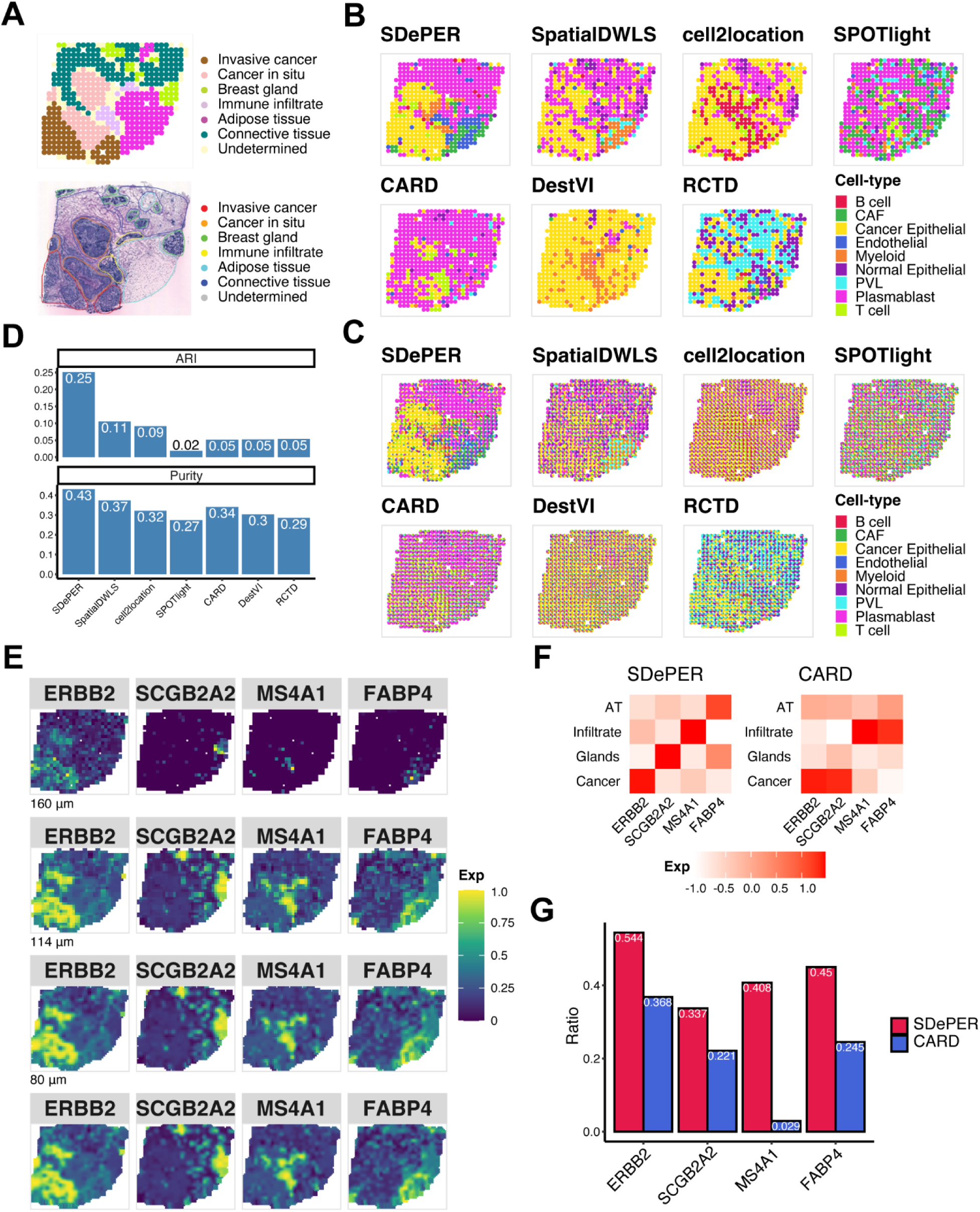
Performance evaluation and comparison using breast cancer dataset. (A) H&E staining of breast cancer and annotated regions. (B) Visualization of inferred dominant cell type in each spot (CAF: cancer-associated fibroblasts; PVL: perivascular-like). (C) Spatial scatter pie chart of estimated cell type composition within each spot. (D) Comparing deconvolution methods using ARI (left) and purity (right). (E) Expression patterns of the corresponding region-specific marker genes and its imputed expression at three different resolution levels: 160μm (about 64% of original size), 114μm (about 32% of original size), 80μm (about 16% of original size). (F) Heatmap showing average imputed expression of region-specific marker genes at 80μm level within each annotated region for SDePER and CARD (AT: adipose tissue; Infiltrate: immune infiltrate; Glands: breast glands; Cancer: invasive cancer and cancer *in situ*). (G) Bar plot showing the ratio of average layer-specific marker gene expression in the corresponding layer among all layers.

The SDePER predicted dominant cell type had the best resemblance to the boundaries between tumor and normal regions in the H&E staining image (Fig. 5B), while other methods failed to detect regions annotated in the staining image. SPOTlight failed to detect any cancer epithelial in the cancer regions, whereas DestVI and cell2location predicted almost all spots to be mainly cancer epithelial cells, which was inconsistent with the H&E staining image. RCTD, SpatialDWLS and CARD had vague boundary between tumor and normal regions. SDePER also achieved the highest ARI and purity, which were 1.24 and 2.36 times higher than the second-best method, respectively, confirming the best performance of SDePER (Fig. 5D). The least non-specific cell types were detected by SDePER (Fig. 5B-C).

For the imputation results, we identified 4 region-specific markers from the ST data for the four annotated regions: cancer region, breast glands, immune infiltrate and adipose tissue. The imputed expression map of each gene by SDePER showed more refined and accurate boundary for its corresponding region (Fig. 5E, S10-11). Compared to CARD, the SDePER imputed expression of each region marker gene separated its corresponding region from the other regions better demonstrated both by visualization (Fig. 5F) and quantitative measure (Fig. 5G). Furthermore, imputed expression of *ERBB2* showed an increase in the cancer region matching previous literature [39]. The expression map of known plasma cell marker gene (Fig. S9), *IGKC*, also ascertained the prevalence of plasma cells in breast glands and connective tissue as predicted by SDePER, rather than perivascular-like (PVL) cells predicted by RCTD. This was also confirmed by the original paper[8]. These further confirmed the higher accuracy of imputation by SDePER.

### Idiopathic pulmonary fibrosis lung data

Lastly, we generated the ST data of a frozen human explant lung sample with Idiopathic Pulmonary Fibrosis (IPF), using the 10x Genomics Visium platform. IPF is a progressive and irreversible, scarring and fibrotic lung disease that leads to a complete remodeling of the lung architecture. Due to the complexity and distortion of lung architecture in fibrotic frozen tissues, only respiratory airway and blood vessels were confidently annotated by a lung fibrosis expert pathologist (Fig. 6A). For deconvolution, we utilized the scRNA-seq dataset of IPF distal lung parenchyma samples as the reference data [40].

**Figure 6.**
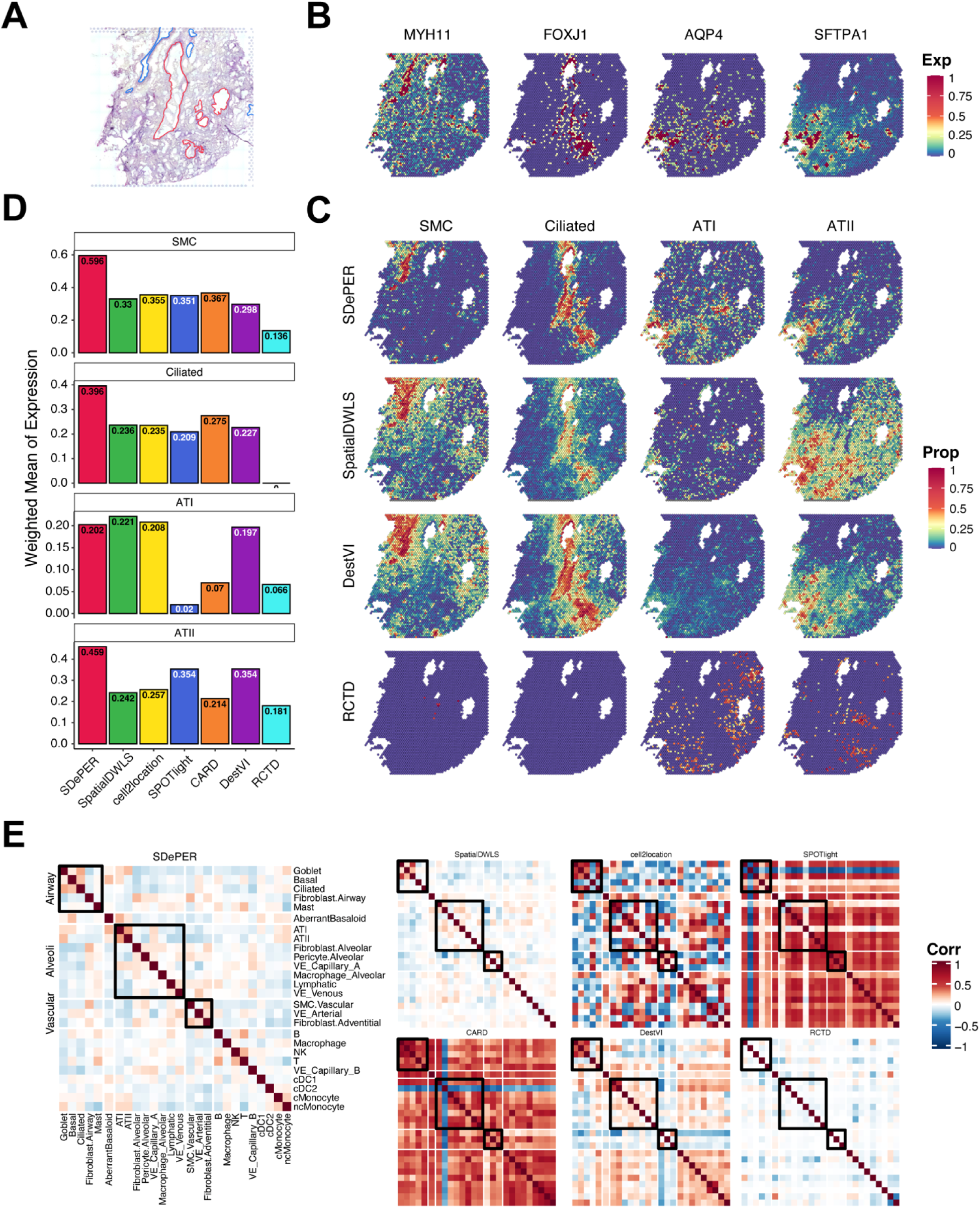
Performance evaluation and comparison using idiopathic pulmonary fibrosis lung dataset. (A) H&E staining of breast cancer with annotated regions: respiratory airway (red) and blood vessels (blue). (B) Heatmaps of selected cell-type marker genes expression patterns for SMC (MYH11), Ciliated cells (FOXJ1), AT1 (AQP4) and AT2 (SFTPA1) cells. (C) Heatmaps of estimated cell-type proportions for SMC, Ciliated cells, AT1 and AT2 cells from SDePER, SpatialDWLS, DestVI and RCTD. (D) Barplot of the average expression of marker genes among all spots weighted by estimated proportions of the corresponding cell type for each method. (E) Pairwise correlation of estimated cell-type proportions for each method.

We demonstrated the results using four cell types: ciliated cells from airway, smooth muscle cells (SMC) from vascular, alveolar type 1 (AT1) and type 2 (AT2) cells from alveoli. Expression maps of marker genes for ciliated cells and SMC match the annotations of airway and vascular, respectively (Fig. 6B). Marker gene expression maps of AT1 and AT2 cells also suggested their prevalence over the distal side of the lung where alveoli are located.

Visualization of the predicted cell type proportions (Fig. 6C, S12) showed that SDePER captured the location of four cell types accurately and precisely, which well-matched both the expression map of marker genes (Fig. 6B) and the pathological annotation (Fig. 6A). But other methods either lacked the specificity in estimation or failed to identify cell types. SpatialDWLS and DestVI had excessive non-zero estimations in spots lacking the corresponding marker gene expression, especially for SMC in the vascular region and ciliated cells in the airway. RCTD could not identify the selected cell types at all. For each method, the average expression of each marker gene across all spots weighted by predicted proportion of its corresponding cell type was calculated to quantitatively measure the consistency between the estimated cell type compositions and marker gene expression maps. SDePER achieved the highest weighted mean for SMC, Ciliated cells and AT2 cells (Fig. 6D). It had a comparable performance in AT1 cells. These quantitatively confirmed the highest estimation accuracy of SDePER. Moreover, SDePER results demonstrated co-localization of AT1 and AT2 cells on the margin of tissue slide with the highest pairwise correlation of estimated cell type proportions, which is consistent with the cell type marker gene expression map and anatomy of human lungs (Fig. 6E).

Overall, this is the first time that ST data from human lung sample is used for demonstration of cell type deconvolution. The results showed that SDePER is a reliable method for complex tissue samples with vague structure and rare cell types.

## Discussion

We have developed a novel deconvolution method, SDePER, to deconvolute the spatial barcoding-based transcriptomic data using reference scRNA-seq data, with considerations of platform effects, sparsity and spatial correlation. Through simulations, we demonstrated the superior performance and robustness of SDePER to platform effects and mismatching cell types between ST and reference data. Applications to datasets from various tissue types, species and platforms also showed a superior accuracy in the estimated cell type compositions and imputed gene expression of SDePER.

There are several directions of extensions of SDePER. First, SDePER removes the systematic differences between the ST and scRNA-seq data in an unsupervised manner, which can be improved by utilizing the known cell type compositions of reference scRNA-seq data and pseudo-spot data as supervision in the training of CVAE. Multi-task learning strategy can be used to integrate the unsupervised and supervised learning and leverage the information of cell type compositions in the pseudo-spot data to guide the CVAE training. Second, we assumed that the distribution of embeddings in the CVAE latent space follows a standard normal distribution. This assumption can be relieved by introducing importance sampling [41, 42]. Third, the encoder and decoder in CVAE are multi-layer neural networks, which are generic to approximate any functions [43, 44], leading to its relative high variance and sensitivity to the variation of model structure and initialization. This can be improved by using negative binomial distribution [45] instead in the decoder. Finally, the computational efficiency of SDePER can be improved by caching the calculated log-likelihoods in GLRM fitting to avoid repetitive.

## Methods

### SDePER method overview

SDePER is built upon the combination of a conditional variational autoencoder (CVAE) [33] and a graph Laplacian regularized regression model (GLRM). The CVAE component aims to remove platform effects and the GLRM component aims to estimate cell type compositions at each spot based on cell type-specific signatures from reference scRNA-seq data with considerations of sparsity and spatial correlation of cell type compositions between neighboring spots in the tissue. Based on the estimated cell type proportions, imputation of cell type compositions and gene expression at unmeasured locations in refined spatial maps with higher resolution is performed using a nearest neighbor random walk.

### Conditional variational autoencoder for platform effect adjustment

The CVAE model considers the two technology platforms, scRNA-seq and ST, as two conditions. To form the training dataset, pseudo-spot data are generated from the reference scRNA-seq data to provide a wide spectrum of cell type compositions for the input under the scRNA-seq condition. For each pseudo-spot, we randomly select a set of cells from reference scRNA-seq data and calculate the average normalized gene expression across cells as the expression profile for the pseudo-spot. The range of the number of selected cells per pseudo-spot is specified based on the cell density in real ST data. In total, the number of pseudo-spots generated is min (100 × *N* × *K*, 500,000), where *N* is the number of spots in the real ST data and *K* is the number of cell types in the reference scRNA-seq data. We train the CVAE model using 80% of the pseudo spots, the reference scRNA-seq data and real ST data. The rest 20% pseudo-spots are used as validation data for learning rate decay and early stopping. Genes used in CVAE are the union of top highly variable genes and cell type marker genes identified from the reference scRNA-seq data identified using Scanpy 1.9.1 [46]. The sizes of both gene lists can be tuned by users based on the properties of reference scRNA-seq data. The normalized expression of each gene is further rescaled separately for the scRNA-seq and ST condition to be from 0 to 10 using min-max scaling. The conditional variable in CVAE is set to 10 for the ST condition (real ST data) and 0 for the scRNA-seq condition (pseudo-spot data and reference scRNA-seq data).

In the CVAE training, we set the number of neurons in latent space as 3 times the number of cell types in the reference scRNA-seq data and use 1 hidden layer for both encoder and decoder under each condition, in which the number of neurons is the largest integer no more than the geometric mean of the number of neurons in the input layer and latent space. We use Adam [47] for optimization and the initial learning rate is set to 0.003 with decay specified based on the value of loss function of the validation dataset. The number of epochs is set to 1,000 and the early training stopping criteria is when the value of loss function of the validation dataset increases. After the CVAE training, the real ST data is embedded into the latent space for the ST condition (conditional variable=10), and then is decoded using the decoder for the scRNA-seq condition (conditional variable=0). The reference scRNA-seq data is encoded and decoded for denoising using the encoder and decoder for the scRNA-seq condition (conditional variable=0). The decoded gene expression levels are min-max scaled back using the scRNA-seq rescaling factors. For the real ST data, the rescaled values are further multiplied by 10,000 and rounded to nearest integers. By using the same decoder, ST and scRNA-seq data are transformed to the same space to remove platform effects. The transformed real ST data and reference scRNA-seq data serve as input to the GLRM component.

### Graph Laplacian regularized model for cell type deconvolution

We fit a graph Laplacian regularized model to estimate cell type compositions in each spot using the transformed ST and reference scRNA-seq data. Only cell type marker genes identified from the transformed reference scRNA-seq data are included in the model. The transformed ST data of each gene is assumed to follow a Poisson distribution with the log-transformed mean being a linear combination of its transformed across all cell types, which forms the base model. The transformed expression profile of each cell type is calculated as the average expression profiles across cells of the given cell type from the transformed reference scRNA-seq data. On top of the base model, we incorporate the spot location information using graph Laplacian regularization that encourages cell type compositions of neighboring spots to be similar. We also enforce cell type sparsity within each spot using adaptive LASSO regularization. Specifically, the GLRM component consists of the following three major components.

#### Base model

The transformed ST data is modeled using a Poisson-loglinear model. For each spot *i* and gene *j*, the transformed ST count *Y_ij_* is assumed to follow a Poisson distribution:

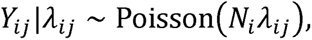

where *N_i_* represents the total UMI count of spot *i* and *λ_ij_* represents the true underlying relative expression level of gene *j* in spot *i*. The rate parameter *λ_ij_* is further modeled as a combination of expression profiles of all *K* cell types weighted by the cell type proportions,

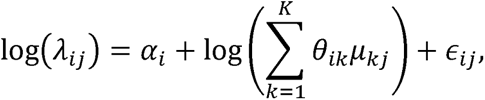

where *α_i_* is a spot-specific fixed effect, *θ_ik_* is the proportion of cells from cell type *k* in spot *i*, *μ_kj_* is the mean expression level of gene *j* in cell type *k* calculated from the transformed reference scRNA-seq data, and *ϵ_ij_* is a random error that follows a normal distribution with mean 0 and variance *σ*^2^ as defined in RCTD[12]. The distribution of *ϵ_ij_* is relaxed to include a heavy tail using an approximation to a Cauchy–Gaussian mixture distribution, which is robust to outliers[48],

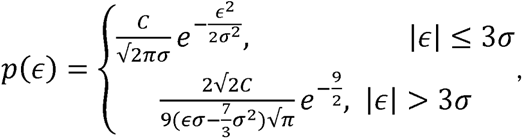

where *C* is a normalizing constant which is chosen to make *p*(*ϵ*) integrate to 1. The model parameters *θ_ik_* are subject to the constraint that 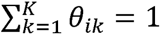 and *θ_ik_* ≥ 0 for all *i* and *k*.

#### Adaptive LASSO regularization

To enforce the local sparsity of cell types in each spot, we penalize the likelihood function of base model using the adaptive Lasso penalty [49]. For each spot *i*, we define the adaptive Lasso penalty as

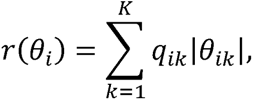

where *q_ik_* is the reciprocal of MLE *θ_ik_* from base model, serving as the weight of *θ_ik_* in the adaptive Lasso penalty.

#### Laplacian regularization

To incorporate spatial information, we represent the physical proximity between spots using an adjacency matrix ***A*** = [*A_ij_*]_*I*×*I*_, where *A_ij_* is an indicator of whether spot *i* and *j* are neighbors on the tissue slide, and *I* is the total number of spots. The graph Laplacian is defined as ***L*** = ***D*** – ***A***, where ***D*** is a diagonal matrix with *D_ii_* = ∑_*j*_*A_ij_*, the degree of spot *i*. The graph Laplacian penalty is defined as

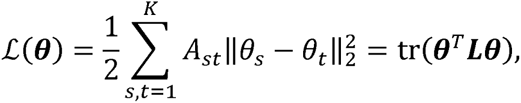

where tr(·) is the trace of a matrix. This penalty measures the aggregate deviation of *θ* between neighboring spots and therefore encourages *θ* to be similar across neighboring spots.

Taken together, we fit the GLRM model by minimizing the following objective function

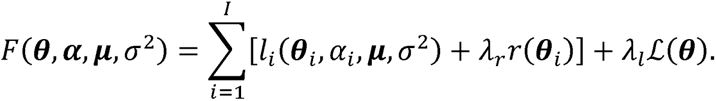

The first term of the objective function is the negative log-likelihood of the base model. The second term is the local adaptive Lasso penalty that enforces cell types to be sparsely present within each spot. The third term is the graph Laplacian penalty to encourage smoothness of cell type compositions across neighboring spots. *λ_r_* and *λ_l_* are two positive hyper-parameters.

The objective function is minimized using a two-stage strategy. To provide initial values of all parameters (***θ***, ***α*** and *σ*^2^) for optimization, we calculate the gene expression profile of cell type *k* using the average library size normalized expression levels of all identified cell type marker genes across all cells of type *k*, denoted as 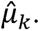 The maximum likelihood estimations 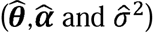 of the base model are obtained using the L-BFGS algorithm [50] in SciPy 1.8.1[51] and serve as the initial values for optimization. In the first stage of optimization, we perform cell type selection in each spot by minimizing the negative log likelihood function of base model with the adaptive LASSO penalty: 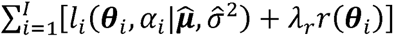 using alternating direction method of multipliers (ADMM) [52]. A cutoff value of 0.001 is applied to the estimate 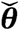 to determine which cell types are present in each spot. In the second stage of optimization, we only include cell types selected from the first stage for each spot in the base model and minimize the negative log likelihood function of base model with graph Laplacian regularization using 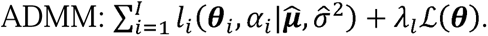 The values of the two hyper-parameters, λ_r_ and *λ_l_*, are chosen using 5-fold cross validation.

### Imputation on cell type composition and gene expression

SDePER borrows information from neighboring spots to perform imputation of cell type compositions at unmeasured locations in refined tissue map with arbitrary resolution by taking the nearest neighbor random walk on the spatial graph. We first use the “finding contour” function in opencv [53] to determine the contours of the tissue shape and distinguish outlines of the tissue and holes inside the tissue. The spatial spots on the outline and border of holes are set to be edge spots. We also develop a custom algorithm to calculate the missing spots in the holes. Suppose the spatial coordinates of the center for spot *i* is *c_i_* = (*x_i_*, *y_i_*) and the distance between centers of two neighboring spots is D. To construct a new spatial map at enhanced resolution, we first create the smallest rectangular region that covers all centers of original spots using [min_*i*_(*x_i_*), max_*i*_(*x_i_*)] × [min_*i*_(*y_i_*), max_*i*_(*y_i_*)]. This rectangular is then gridded into squares with side length *d*(*d* < *D*). Among all the squares, those with center located at *c_i*_* = (*x*_*i*\*_, *y*_*i*\*_) satisfying one of the following criteria are considered as spots in the new map and imputed:

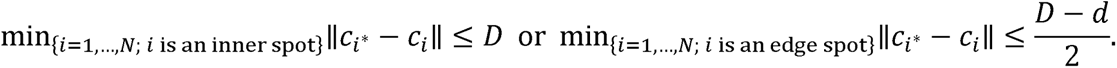

The new spatial map at enhanced resolution (square side length = *d*) consists of the spots with centers ***C***\* = {*c_i*_*}, *i\** = 1, …, *N*\*.

Let ***θ***_*i*\*_ denote the cell type proportions in spot *i*\* in the new spatial map. To perform imputation, we first assign an initial value for ***θ***_*i*\*_ by finding the nearest original spot(s) of spot *i*\* and set 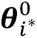 as the average cell type proportion among its neighbors. We construct a Gaussian kernel ***W*** as follows

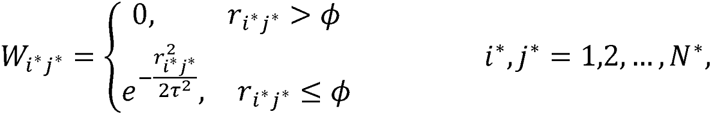

where *r_i*j*_* = ∥*c_i*_* - *c_j*_*∥ is the distance between two spots *i*\* and *j*\* in the new map, *ϕ* is a predefined neighborhood size within which spots contribute to the imputation of each other, and *τ*^2^ is the variance. A nearest neighbor random walk matrix is constructed as ***M*** = ***D*^-1^*W***, where ***D*** is a diagonal matrix with *D_i*j*_* = ∑_*j*\*_*W_i*j*_*. The imputed cell type compositions are obtained by taking a one-step nearest neighbor random walk with the graph which can be written as

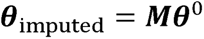

We further impute gene expression at enhanced resolution as ***X***_imputed_ = ***θ***_imputed_***θ*^+^*X***, where ***X*** is the observed UMI counts from ST data, normalized by spot’s sequence depth, and ***θ*^+^** is the Moore-Penrose inverse of ***θ*** with ***θ^+^*** = (***θ***^*T*^***θ***)^-1^***θ***^*T*^.

The hyperparameters in the Gaussian kernel ***W*** include *ϕ* and *τ*^2^. For a given real ST data, the hyperparameter tunning was conducted using a coarse-grained data generated similarly as the simulation study at the resolution the same as the real ST data. Imputation is performed at different values of the hyperparameters and the root mean square error (RMSE) is calculated between the imputed and ground-truth cell type compositions. The hyperparameter value achieving the smallest RMSE is chosen to determine the Gaussian kernel matrix.

### Other deconvolution methods for comparison

Six state-to-art spatial deconvolution methods were chosen to compare with SDePER, including SpatialDWLS (implemented in the R package Giotto, version 1.1.1)[26], SPOTlight (version 1.0.0)[25], RCTD (version 2.0.1)[12], cell2location (version 0.1)[15], DestVI (implemented in the python package scVI, version 0.17.1)[14], and CARD (version 1.0)[24]. We followed the tutorial on the GitHub repository of each method and used the recommended default parameter settings for the deconvolution analyses conducted in this article. When parameters are required to be set manually, we used the values suggested in the vignettes.

### Simulation studies

To evaluate the method performance and demonstrate the impact of platform effects on all deconvolution methods, we simulate spot-level ST data based on the adult mouse primary visual cortex STARmap data that has single-cell resolution [34]. We extract experiments “20180410-BY3_1kgenes” and “20180505_BY3_1kgenes”, and manually put them in the same spatial map with enough space in between so that cells or simulated spots from different experiments are not considered as spatial neighbors. To simulate the ST data, we gridded the tissue slide into squares with side length of ∼□51.5□μm as capture spots, which generated 581 spots with an average of 3.6 cells present per spot. We only kept cells from cell types present in both STARmap data and external reference scRNA-seq data. In each spot, the proportion of cells from each cell type is calculated and serves as ground-truth for performance evaluation. The simulated gene expression level of gene *j* for a given spot that contains cells *i* = 1, …, *n* is calculated as 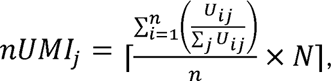 where *U_ij_* is the number of UMIs of gene *j* in cell *i* from the STARmap data and *N* is a fixed scaling factor set to be 1000.

To demonstrate the impact of platform effects on the method performance, each method was applied to the simulated data using two different reference scRNA-seq datasets: internal reference and external reference. The internal reference data is the original STARmap data so there were no platform effects. The external reference data is an independent publicly available adult mouse visual cortex scRNA-seq dataset (GEO accession number: GSE115746) [54]. Under this case, significant platform effects were expected to exist because the simulated ST data and reference data were generated using the *in situ* sequencing and SMART-seq technologies, respectively. We selected 12 overlapping cell types between the external reference data and the STARmap data for deconvolution, which include astrocytes, excitatory neurons layer 2/3, excitatory neurons layer 4, excitatory neurons layer 5, excitatory neurons layer 6, endothelial, microglia, oligodendrocyte, *Pvalb*-positive cells, *Vip* inhibitory neurons, *Sst* neurons and smooth muscle cells. In total, 2,002 cells and 1,020 genes were included in the STARmap data while 11,835 cells and 45,768 genes were in the external reference data.

Deconvolutions using the overlapping 12 cell types between the STARmap data and external reference scRNA-seq data represents analysis scenario 1, under which cells types in the reference data match perfectly with those in the ST data. However, in practice, they can have mismatching cell types, so we modified the external reference data to demonstrate the robustness of all methods to mismatching cell types. In analysis scenario 2, we remove *Vip* inhibitory neurons (n = 1,690) from the reference data. In analysis scenario 3, we added “high intronic” cells (n=182), which are not present in the STARmap data, to the reference data.

To evaluate the method performance, we compare the cell type compositions estimated by each method to the ground-truth across all spots using the root mean square error (RMSE) that quantifies the overall estimation accuracy, Jensen-Shannon Divergence (JSD) that assesses similarity between the estimated cell type distribution and ground-truth per spot, Pearson’s correlation coefficient that measures the similarity of estimation to ground-truth and false discovery rate (FDR) that measures how many cell types were falsely predicted to be present. For the methods without cell type selection procedure, including cell2location, DestVI and CARD, we put a hard-thresholding with cutoff 0.1 on the estimated cell type compositions to force the extremely small proportions to be zeros.

### Real dataset analysis

#### Mouse olfactory bulb dataset

We obtain the mouse olfactory bulb (MOB) ST data from the Spatial Research lab [4]. We focus on the “MOB replicate 12” file which contains 16,034 genes and 282 spots. An independent scRNA-seq data is also downloaded as the reference scRNA-seq data (GEO accession number: GSE121891) [35], which consists of 18,560 genes and 12,801 cells from 5 cell types: granule cells, olfactory sensory neurons, periglomerular cells, mitral and tufted cells, and external plexiform layer interneurons.

To apply SDePER on the MOB dataset, we select 250 highly variable genes and 244 cell type marker genes from the reference scRNA-seq data, which form a set of 434 unique genes used in the CVAE component for platform effects removal. We randomly select 10-40 single cells from the scRNA-seq data to generate a pseudo spot to mimic the number of cells per spot hyper-parameter *λ_r_* is chosen to be 1.931 and *λ_l_* to be 5.179 based on cross-validation. The suggested in the MOB data. 168 cell type marker genes are used in GLRM component. The running time of SDePER is 0.64 hours in total using a 20-core, 100 GB RAM, Intel Xeon 2.6 GHz CPU machine.

#### Melanoma dataset

We download the melanoma dataset from the Spatial Research lab [7]. We focus on the second replicate from biopsy 1 because it contains regions annotated as lymphoid tissue and is extensively examined in the original paper. Biopsy 1 contains 16,148 genes and 293 spots. The reference scRNA-seq dataset is downloaded from GEO database (accession number: GSE115978), which contains 23,686 genes and 2,495 cells from 8 selected samples. In total, 7 cell types are present in the reference data, including malignant cells, T cells, B cells, natural killer (NK) cells, macrophages, cancer-associated fibroblasts (CAFs), and endothelial cells[37].

We choose the top 300 highly variable genes and 280 marker genes, corresponding to 534 unique genes for the CVAE component in SDePER. The number of cells per pseudo-spot is set to be 5-40 cells as provided in the original paper. 145 cell type marker genes are used in GLRM component. The hyper-parameter •_"_ is chosen to be 1.931 and *λ_l_* to be 37.276 based on cross-validation. The running time of CVAE-GLRM is 0.67 hours in total using a 32-core, 100 GB RAM, Intel Xeon 2.6 GHz CPU machine.

#### Breast cancer dataset

We obtain the HER2-positive Breast Cancer spatial transcriptomics dataset from a previous study [8]. The first section of patient H with 15,029 genes and 613 spots is selected for the analysis. We obtain the scRNA-seq data of five HER2-positive tumors from GEO database (accession number: GSE176078) [38] as the reference scRNA-seq data for deconvolution. The reference data consists of 29,733 genes and 19,311 cells from 9 cell types.

For the CVAE component of SDePER, we select the top 1,500 highly variable genes and 824 cell type marker genes, corresponding to 1,942 unique genes. Each pseudo-spot is assumed to contain 20-70 cells based on estimation by other cancer ST studies using the same ST platform [55]. 290 cell type marker genes are used in GLRM component. The hyper-parameter *λ_r_* is chosen to be 1.931 and *λ_l_* to be 37.276 based on cross-validation. The running time of CVAE-GLRM is 2.58 hours in total using a 32-core, 100 GB RAM, Intel Xeon 2.6 GHz CPU machine.

#### Idiopathic pulmonary fibrotic lung dataset

We measured the ST data of a human IPF lung sample from an explanted lung of patient with end stage IPF explanted lung (Yale IRB:1601017047) using the 10x Genomics Visium platform, which is a complex and challenging sample with vague structure and different lung compartments including the bronchi, vascular, mesenchyme and immune compartment. We selected one block of frozen lung tissue obtained from a patient with Idiopathic Pulmonary Fibrosis (IPF). Sections of 10 μm (fresh frozen samples) were cut from the blocks onto Visium slides (10x Genomics) and processed according to the manufacturer’s protocol Tissue sections were Hematoxylin and Eosin stained and finally imaged (20X) using a scanning microscope (EvosM700, ThermoFischer Scientific). Tissue was permeabilized and mRNAs were hybridized to the barcoded capture probes directly underneath. cDNA synthesis connects the spatial barcode and the captured mRNA. After RT and amplification by PCR, Dual-indexed libraries were prepared as in the 10x Genomics protocol, and sequenced (2 samples/ HiSeq 6000 flow cell) with read lengths 28 bp R1, 10 bp i7 index, 10 bp i5 index, 90 bp R2. Base calls were converted to reads with the software SpaceRanger’s implementation mkfastq (SpaceRanger v1.2.2). Multiple fastq files from the same library and strand were catenated to single files. Read2 files were subjected to two passes of contaminant trimming with cutadapt (v1.17): for the template switch oligo sequence (AAGCAGTGGTATCAACGCAGAGTACATGGG) anchored on the 5′ end and for poly(A) sequences on the 3′ end. Following trimming, read pairs were removed if the read2 was trimmed below 30 bp. Visium libraries were mapped on the human genome (10X-provided GRCh38 reference), using StarSolo (StarSolo v2.7.6a).

This data measured expression of 60,651 genes at 4,992 spatial locations. We filtered out the spatial location not covered by tissue and the genes not expressed on all spatial locations. Finally, we performed cell type deconvolution on 32,078 genes and 3,532 spatial spots. We used the scRNA-seq data from an IPF lung in a previous study as the reference [40]. This dataset consists of 60,651 genes and 12,070 cells. These cells have already been annotated into 39 cell types. Some of the cell types had insufficient cells to provide sufficient information to perform deconvolution. We reannotated the scRNA-seq dataset with 44 cell types in total and selected major cell types with sufficient number of cells. Therefore, we only considered 11,227 cells from 26 major cell types, and we further filtered out the gene not expressed on these cells. Finally, a final set of 35,483 genes and 11,227 cells serves as the reference scRNA-seq data for deconvolution.

We choose the top 2,000 highly variable genes and a set of manually selected 2,534 marker genes, corresponding to 3,101 unique genes for the CVAE component in SDePER. The number of cells per pseudo-spot is set to be 2-10 cells as provided by expert advice. 1,788 cell type marker genes are used in GLRM component. The hyper-parameter *λ_r_* is chosen to be 0.72 and *λ_l_* to be 13.895. The running time of SDePER is 109.27 hours in total using a 64-core, 100 GB RAM, Intel Xeon 2.6 GHz CPU machine.

## Data Availability

This study uses both publicly available datasets and a private dataset generated in the laboratory of Dr. Naftali Kaminski. The publicly available datasets include the STARmap dataset (https://kangaroo-goby.squarespace.com/data), MOB dataset (https://www.spatialresearch.org/resources-published-datasets/doi-10-1126science-aaf2403/), melanoma dataset (https://www.spatialresearch.org/resources-published-datasets/doi-10-1158-0008-5472-can-18-0747/), breast cancer dataset (https://doi.org/10.5281/zenodo.4751624). The IPF dataset has been deposited to the GEO database.

## Code Availability

The SDePER source code and software are free at https://github.com/az7jh2/SDePER under MIT license. We also provide a Docker image of SDePER at https://hub.docker.com/r/az7jh2/sdeper. All scripts used to conduct all the analysis are available at https://github.com/az7jh2/SDePER_Analysis.

## Acknowledgements

This study was supported by the National Institutes of Health (NIH) grants R21LM012884 (to X.Y.), R01LM014087 (to Z.W. and X.Y.), National Science Foundation (NSF) grant DMS1916246 (to Z.W.). A.J. is supported by funding from Fond de dotation du Souffle, Philippe Foundation, Bourse de Mobilite CHU de Caen, Bourse de mobilite internation interregion Nord Ouest G4.

## Authors’ contributions

Z.W. and X.Y. conceived the idea and provided funding support. N.L, Y.L., A.A., Z.W. and X.Y. designed the study. N.L., Y.L., J.Q. and G.X. developed the method, implemented the software and performed simulations. Y.L., N.L., J.Q., J.Z., N.W., X.H., and W.J. conducted the real data analysis. A.J., T.S.A., I.R., R.H. and N.K. provided the IPF dataset. A.J., T.S.A, and N.K. aided result interpretation. Y.L., N.L., J.Q., Z.W. and X.Y. wrote the manuscript. Z.W. and X.Y. supervised the research. All authors read and approved the final manuscript.

