## Supplementary Information for "A hybrid machine learning and regression method for cell type deconvolution of spatial barcoding-based transcriptomic data"

### Supplementary Figures

**Supplementary Figure S1. Comparison of the accuracy in estimated cell type proportions.** The true and predicted cell type proportion of (A) L4 excitatory neurons, (B) L5 excitatory neurons, (C) L6 excitatory neurons, and (D) oligodendrocytes by each method across all simulated spots are shown.

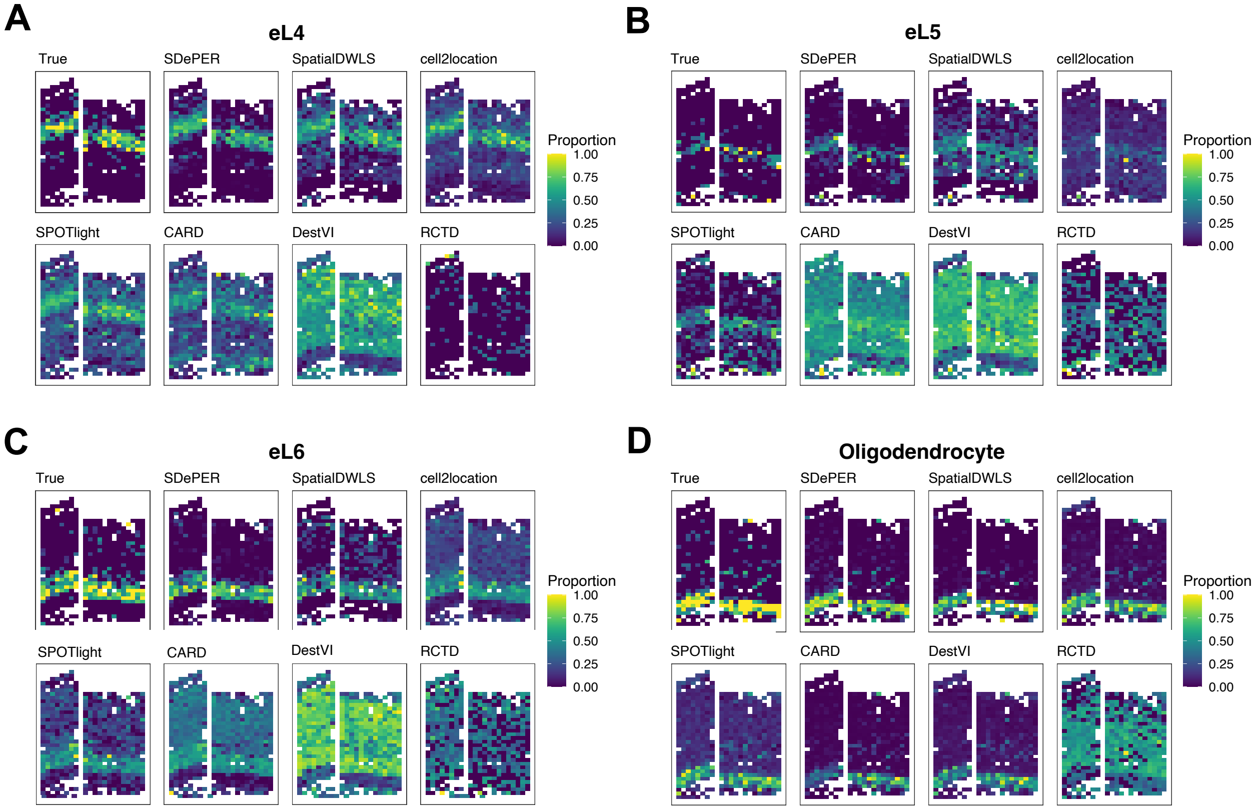

**Supplementary Figure S2. Correlation between predicted cell type proportion and ground-truth in simulated data.** For each method and each type of reference data (external or internal), the correlation matrix between predicted cell type proportion and underlying true proportion is shown.

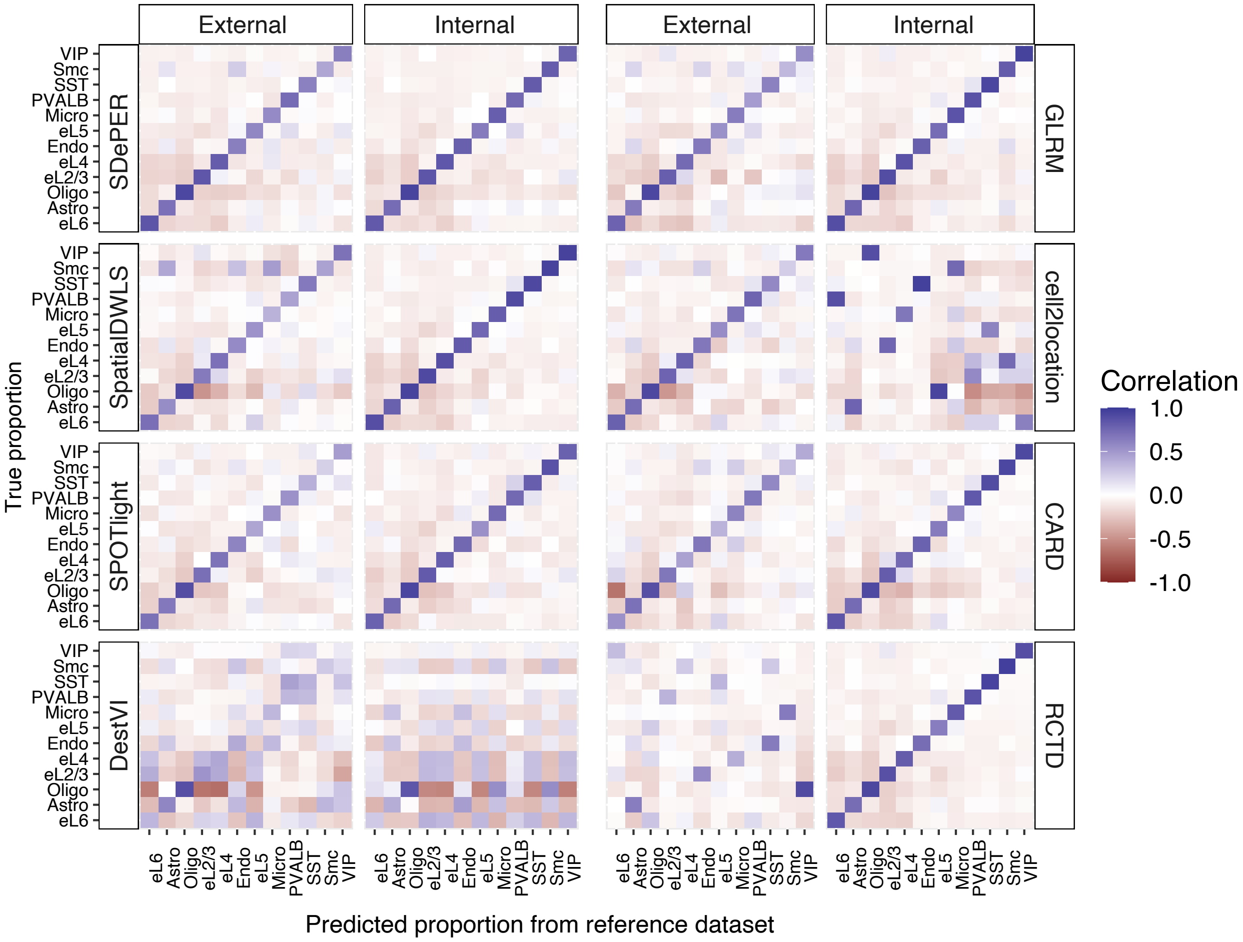

**Supplementary Figure S3. Expression map of marker genes of GC, M/TC, PGC, OSNs, EPL-IN in the MOB dataset.** Comparison of these heatmaps to Fig. 3A confirms the dominance of each cell type in the corresponding annotated layer based on the histology staining.

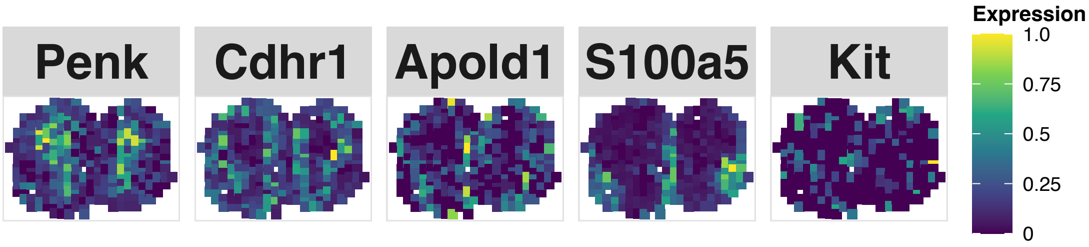

**Supplementary Figure S4. Comparison of the predicted and imputed cell type proportion at different resolution levels between SDePER and CARD in the MOB dataset.** The original resolution was 200 $\mu m$ which correspond to the estimated cell type proportions from the deconvolution. The three higher resolution levels included 160, 114 and 80$\mu m$, for which the cell type proportions were imputed based on the estimated cell type proportions at the original resolution (200 $\mu m$).

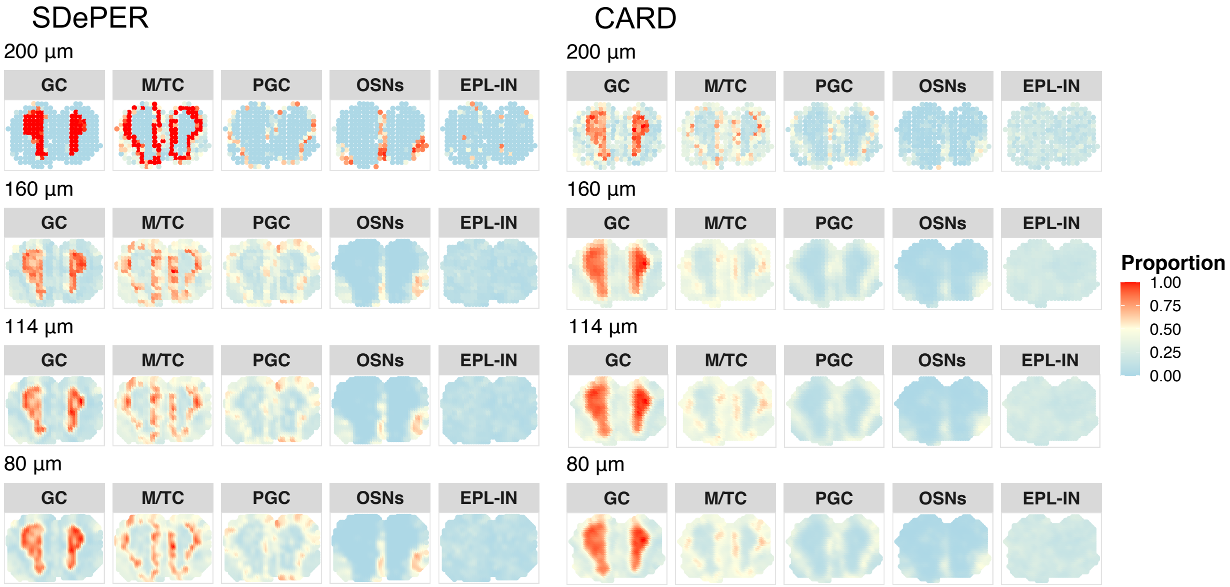

**Supplementary Figure S5. Visualization of the imputed expression of layer marker genes at different resolution levels by SDePER and CARD in the MOB dataset.** Three enhanced resolution levels included 160, 114 and 80$\mu m$, for which the expression of each gene was imputed based on the predicted cell type proportions and the ST data at the original resolution (200 $\mu m$).

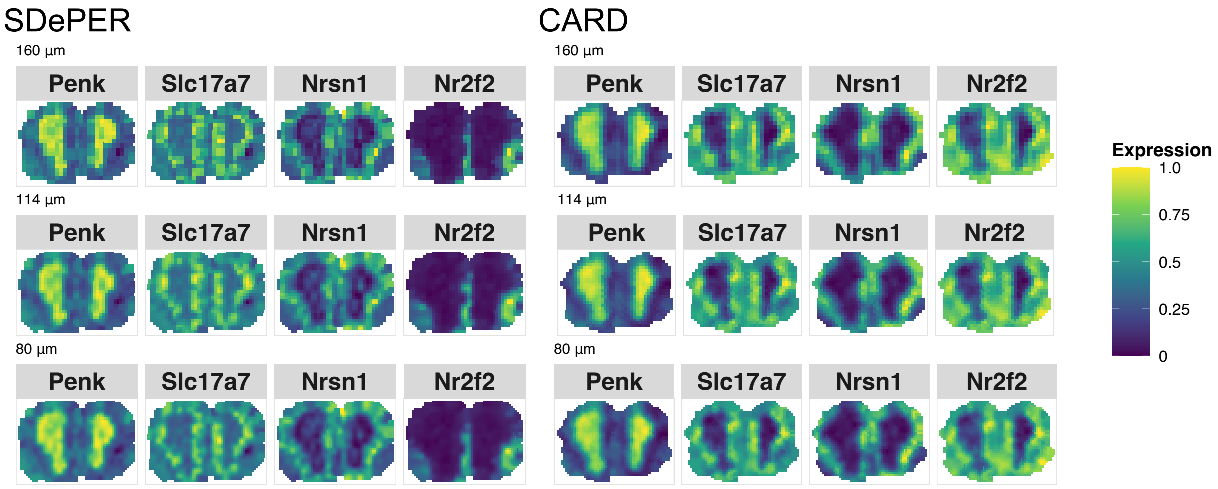

**Supplementary Figure S6. Visualization of expression of cell type marker genes in the melanoma dataset.** One marker was chosen per cell type and demonstrated from left to right and top to bottom for malignant, CAF, macrophage, B cell, T cell, NK cell, endothelial in melanoma spatial dataset, respectively.

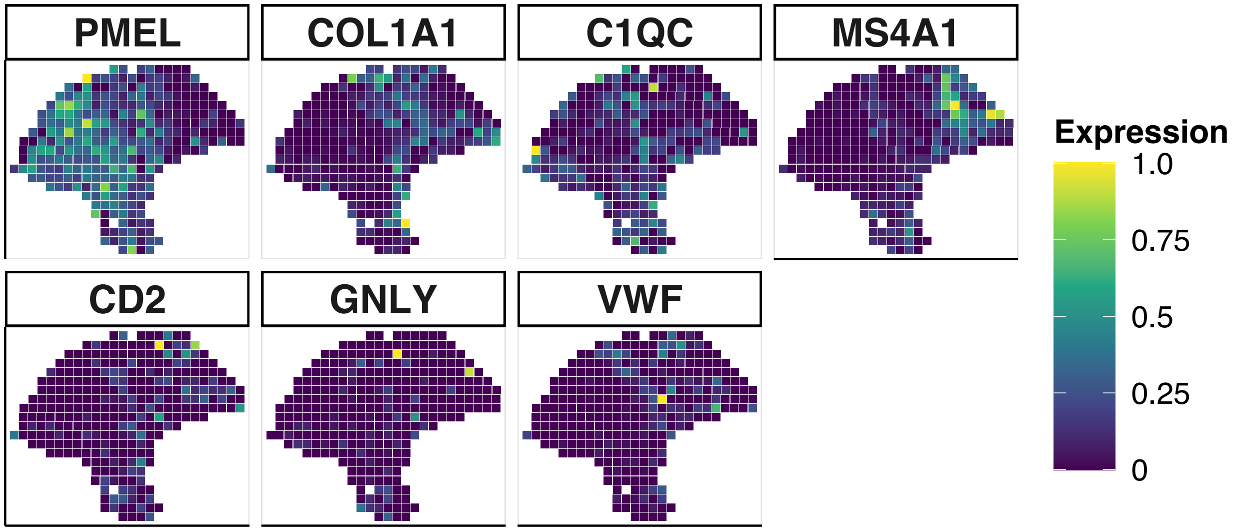

**Supplementary Figure S7. Comparison of the predicted and imputed cell type proportion at different resolution levels between SDePER and CARD in the melanoma dastaset.** The original resolution was 200 $\mu m$ which correspond to the estimated cell type proportions from the deconvolution. The three higher resolution levels included 160, 114 and 80$\mu m$, for which the cell type proportions were imputed based on the estimated cell type proportions at the original resolution (200 $\mu m$).

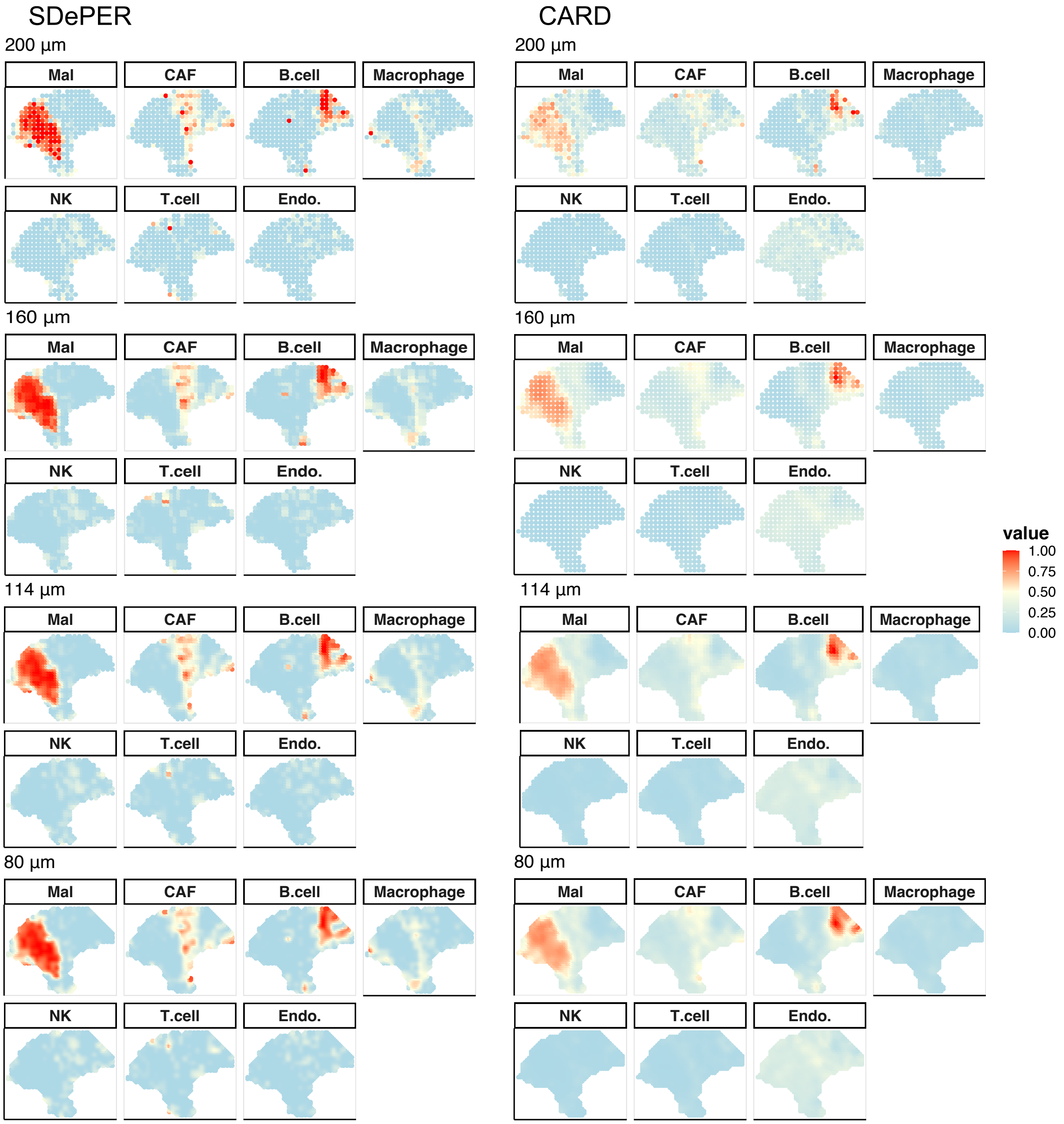

**Supplementary Figure S8. Visualization of the imputed expression of layer marker genes at different resolution levels by SDePER and CARD in the melanoma dataset.** Three enhanced resolution levels included 160, 114 and 80$\mu m$, for which the expression of each gene was imputed based on the predicted cell type proportions and the ST data at the original resolution (200 $\mu m$).

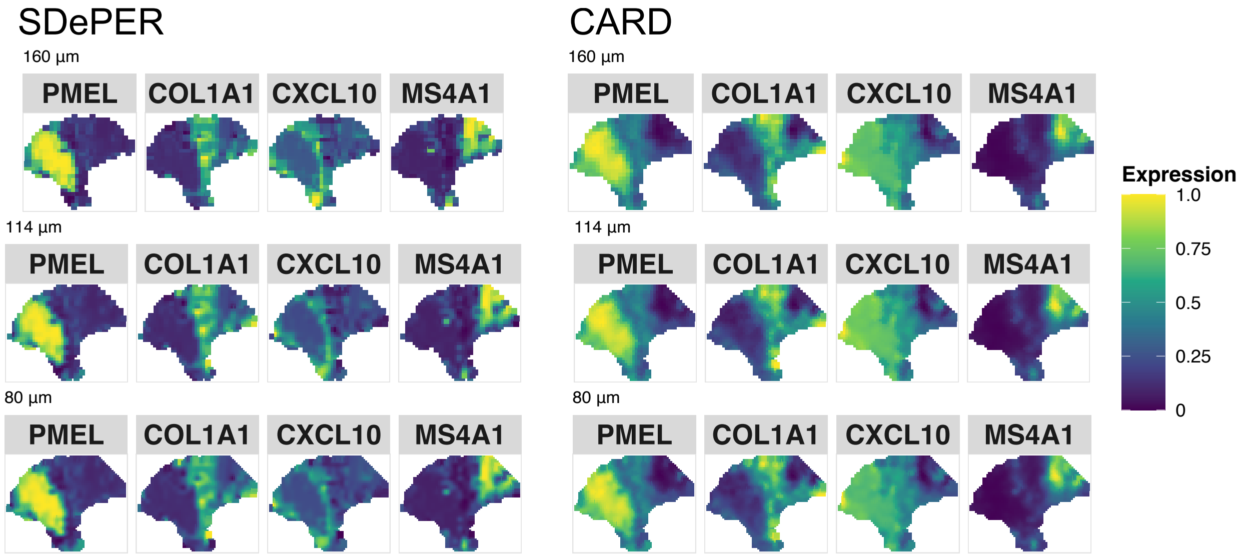

**Supplementary Figure S9. Expression map of marker genes for different cell types in the breast cancer dataset.** One marker was chosen per cell type and demonstrated from left to right and top to bottom for cancer epithelial, CAF, plasma, myeloid, PVL, endothelial, B cell, T cell, normal epithelial, respectively.

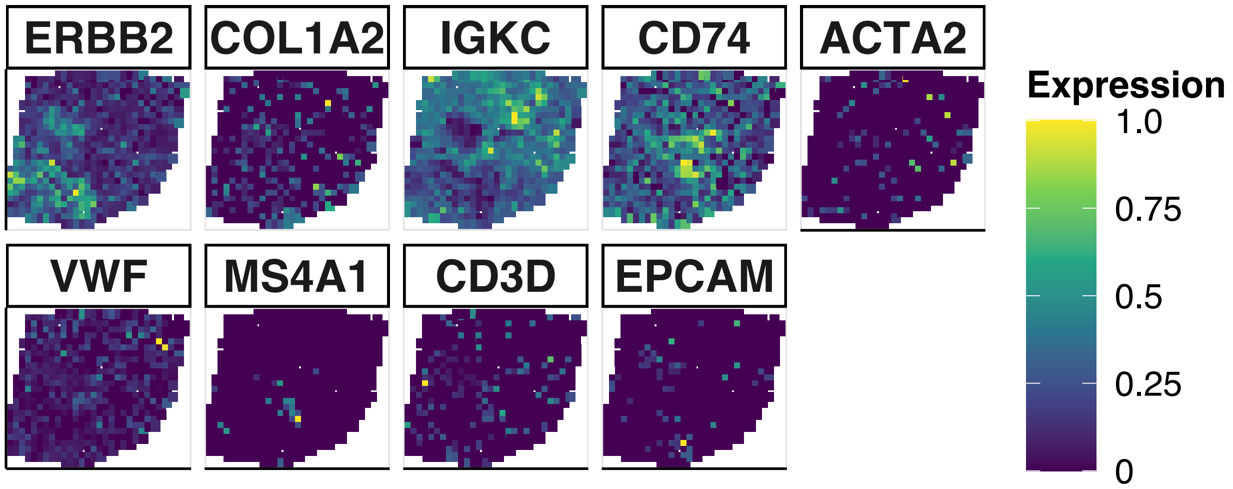

**Supplementary Figure S10. Comparison of the predicted and imputed cell type proportion at different resolution levels between SDePER and CARD in the breast cancer dataset.** The original resolution was 200 $\mu m$ which correspond to the estimated cell type proportions from the deconvolution. The three higher resolution levels included 160, 114 and 80$\mu m$, for which the cell type proportions were imputed based on the estimated cell type proportions at the original resolution (200 $\mu m$).

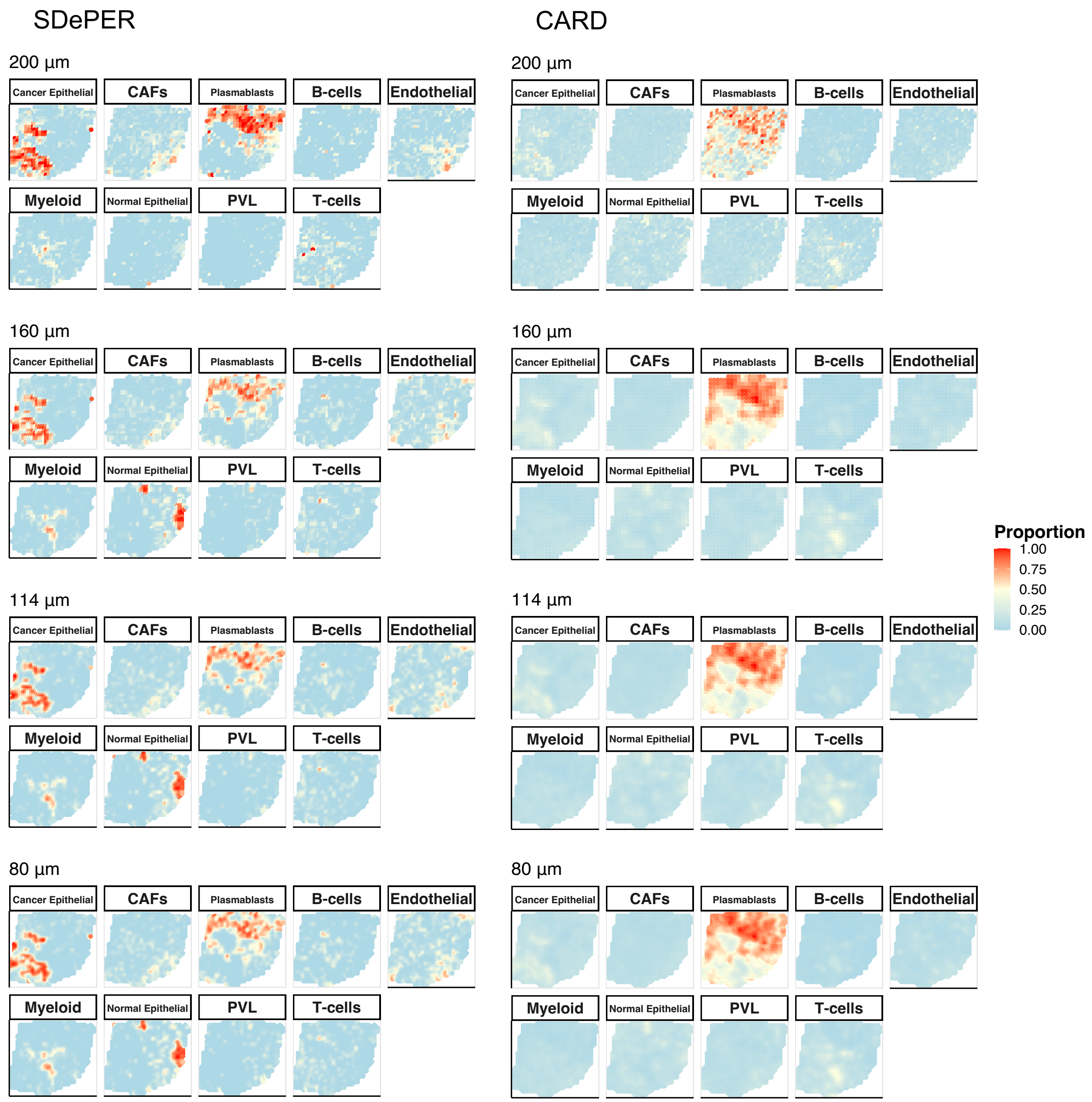

**Supplementary Figure S11. Visualization of the imputed expression of layer marker genes at different resolution levels by SDePER and CARD in the breast cancer dataset.** Three enhanced resolution levels included 160, 114 and 80$\mu m$, for which the expression of each gene was imputed based on the predicted cell type proportions and the ST data at the original resolution (200 $\mu m$).

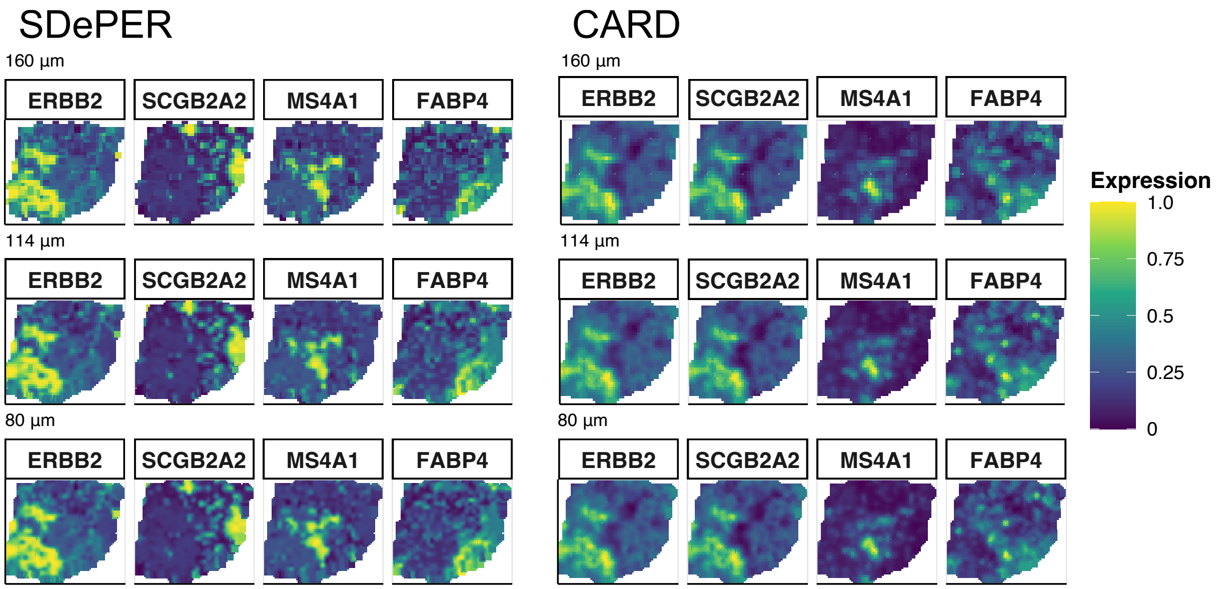

**Supplementary Figure S12. Visualization of the predicted cell type proportions by all methods in the IPF dataset.** Four cell types, including AT1, AT2, SMC and Ciliated cells, were demonstrated from top to bottom, respectively.

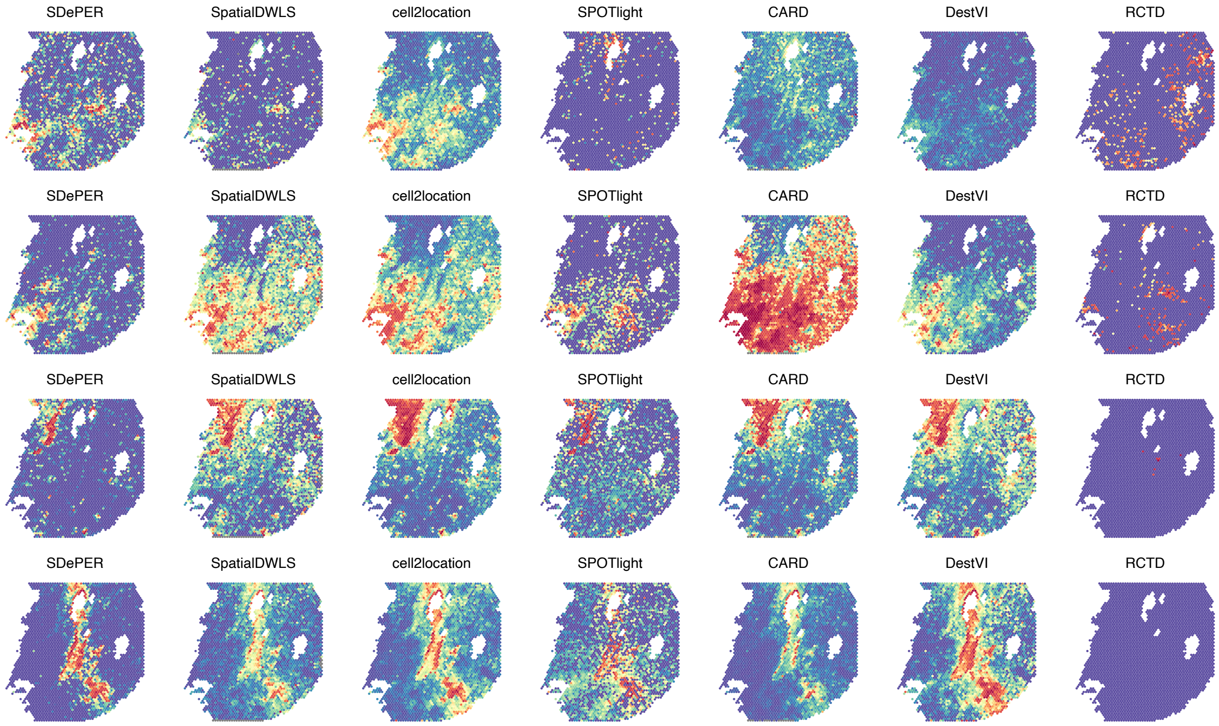

#### Supplementary Tables

**Supplementary Table S1. Quantitative comparison of the accuracy in estimated cell type proportions.** Results for internal and external reference were shown. The median RMSE, JSD, correlation coefficient and FDR was used for the comparison.

| Methods | Reference | median_RMSE | median_JSD | median_cor | median_FDR |
|---|---|---|---|---|---|
| **SDePER** | Internal | 0.102 | 0.191 | 0.888 | 0.500 |
| | External | 0.121 | 0.269 | 0.828 | 0.750 |
| **GLRM** | Internal | 0.081 | 0.129 | 0.928 | 0.500 |
| | External | 0.172 | 0.458 | 0.565 | 0.800 |
| **SpatialDWLS** | Internal | 0.074 | 0.104 | 0.945 | 0.500 |
| | External | 0.153 | 0.397 | 0.658 | 0.750 |
| **cell2location** | Internal | 0.252 | 0.861 | -0.102 | 0.833 |
| | External | 0.171 | 0.478 | 0.635 | 0.833 |
| **SPOTlight** | Internal | 0.151 | 0.384 | 0.821 | 0.818 |
| | External | 0.179 | 0.504 | 0.664 | 0.818 |
| **CARD** | Internal | 0.113 | 0.180 | 0.855 | 0.333 |
| | External | 0.199 | 0.486 | 0.383 | 0.667 |
| **DestVI** | Internal | 0.203 | 0.580 | 0.335 | 0.750 |
| | External | 0.209 | 0.626 | 0.285 | 0.750 |
| **RCTD** | Internal | 0.102 | 0.138 | 0.932 | 0.000 |
| | External | 0.238 | 0.682 | 0.051 | 0.750 |
